# Calculating the force-dependent unbinding rate of biological macromolecular bonds from the force-ramp optical trapping assays

**DOI:** 10.1101/2021.09.20.461030

**Authors:** Apurba Paul, Joshua Alper

**Affiliations:** Department of Physics and Astronomy, Clemson University, Clemson, SC; Eukaryotic Pathogen Innovation Center, Clemson University, Clemson, SC; Department of Biological Sciences, Clemson University, Clemson, SC

## Abstract

The non-covalent biological bonds that constitute protein-protein or protein-ligand interactions play crucial roles in many cellular functions, including mitosis, motility, and cell-cell adhesion. The effect of external force (*F*) on the unbinding rate (*k*_off_(*F*)) of macromolecular interactions is a crucial parameter to understanding the mechanisms behind these functions. Optical tweezer-based single-molecule force spectroscopy is frequently used to obtain quantitative force-dependent dissociation data on slip, catch, and ideal bonds. However, analyses of this data using dissociation time or dissociation force histograms often quantitatively compare bonds without fully characterizing their underlying biophysical properties. Additionally, the results of histogram-based analyses can depend on the rate at which force was applied during the experiment and the experiment’s sensitivity. Here, we present an analytically derived cumulative distribution function-like approach to analyzing force-dependent dissociation force spectroscopy data. We demonstrate the benefits and limitations of the technique using stochastic simulations of various bond types. We show that it can be used to obtain the detachment rate and force sensitivity of biological macromolecular bonds from force spectroscopy experiments by explicitly accounting for loading rate and noisy data. We also discuss the implications of our results on using optical tweezers to collect force-dependent dissociation data.

## Introduction

The weak non-covalent bonds that constitute protein-protein and protein-ligand interactions underlie nearly all cellular functions^1^. For example, the physical chemistry of protein-protein interactions is fundamental to the molecular mechanisms of the cytoskeleton^2^. Kinesin and dynein motor proteins precisely regulate and coordinate changes in motor-filament binding affinity as a function of their mechanochemical cycles as they walk along microtubules, and quantitative models of motors must explicitly account for the effects of external forces (*F*) on the filament unbinding rate constant (*k*_off_(*F*))^2^. Beyond motors, examples of force-dependent unbinding rate constants are ubiquitous in biological systems, including cell-cell adhesion^3–5^, mechanotransduction^6,7^, DNA polymerases and helicases^8,9^, membrane-surface adhesion^10^, selectin-ligand^11^, and antibody-antigen complexes^12–14^.

Weak non-covalent bonds between biological macromolecules can be classified into three principal categories^15^. Slip bonds have a lifetime that decreases with the increasing applied force^16^. Catch bonds have a lifetime that increases with applied force^10,17^. Ideal bonds have a lifetime that is independent of applied force^10^. Many biomolecular interactions have been characterized using single-molecule techniques and understood in the context of these models.

Single-molecule force spectroscopy experiments involving the force-dependent unbinding of motor proteins^18^, microtubule- and actin-associated proteins^19–21^, focal adhesion proteins^22,23^, and many others^24–26^, are commonly done using optical tweezers to apply a constant (i.e., a step function in time) or linearly increasing force (e.g., a force ramp starting a zero upon binding and increasing until detachment) to the molecules (Figure 1a). Data calculated from force spectroscopy traces frequently include the bound time and unbinding force (Figure 1b). For example, recent single-molecule experiments on cytoplasmic dynein motor proteins show both catch bond and slip-ideal bond behaviors^27,28^ and that kinesin exhibits force-dependent stepping velocities and microtubule unbinding rates that are a function of force direction^29^. Multiple techniques exist to analyze this data^30^, including a recent work extending classical analysis techniques to characterize motor proteins pulling cargo from a stationary optical trap^31^.

**Figure 1.**
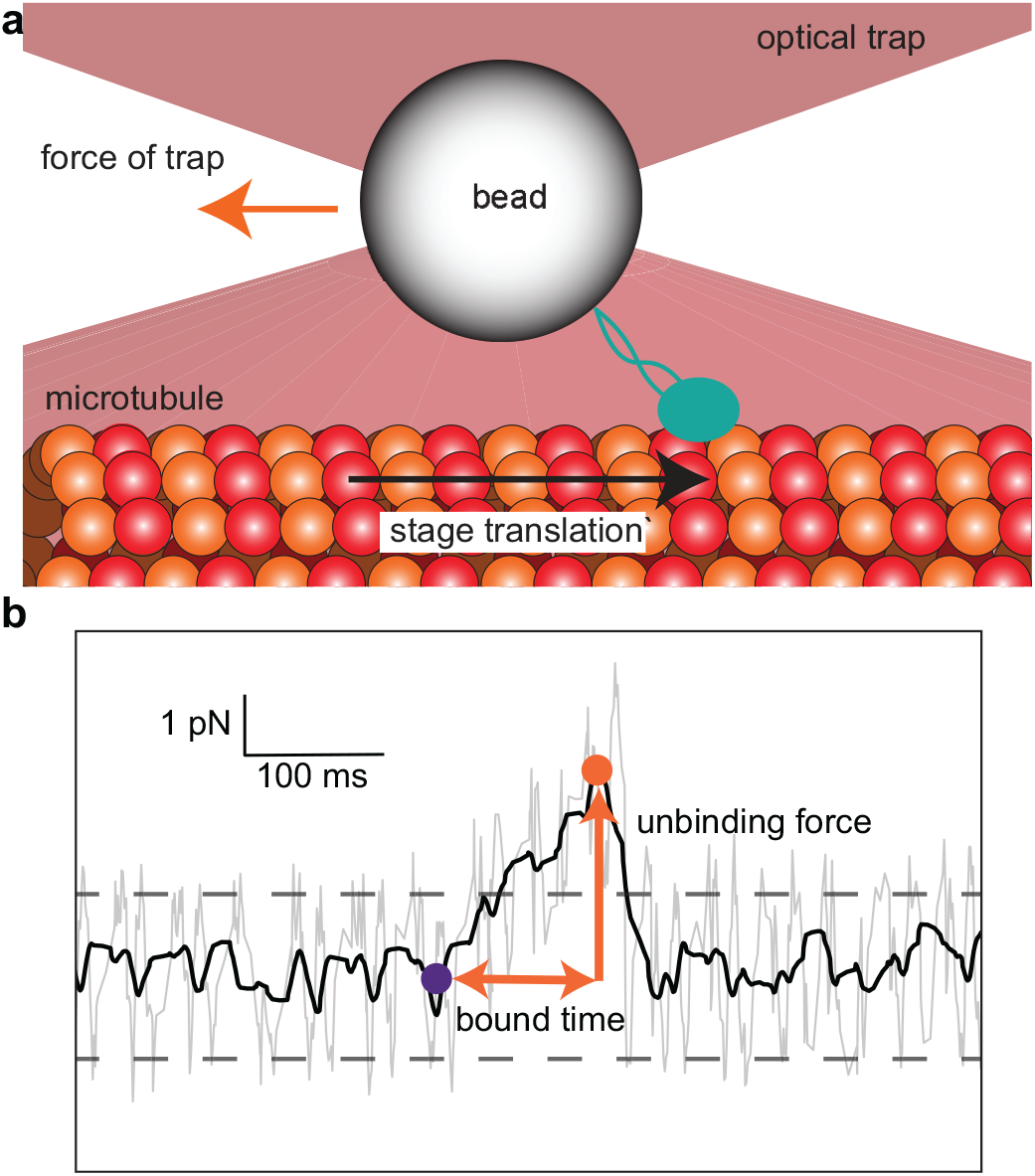
Microtubule-associated protein dissociation from a microtubule in a force-ramp optical trapping assay. **(a)** Schematic of a microtubule (*red* and *orange* circles) bound to a stage that moves (with velocity, *v, black arrow*) and pulls a bead conjugated to a microtubule-associated protein (MAP, *green*) out of the trap center (with effective stiffness of the trap and MAP, *κ*). The force that the trap exerts on the bead (*orange arrow*) increases at a rate of *f* = *κv* (thus *F*(*t*) = *ft*) until the MAP stochastically unbinds from the microtubule, which causes it to return to the trap center. The force-dependent unbinding rate constant, *k*_off_(*F*), governs the biophysics of unbinding. **(b)** A typical force spectroscopy trace (*gray* = raw data, *black* = filtered data) from a force-ramp optical tweezer assay showing the force as a function of time during a biological macromolecular binding (initial binding, *purple dot*) and unbinding (final unbinding, *orange dot*) event. The unbinding force and bound times (*orange arrows*) are used to determine *k*_off_(*F*) for the MAP-microtubule interaction. An event is “detected” when the filtered signal exceeds 5-times the standard deviation of the noise (*dashed lines*).

This report presents a cumulative distribution function approach to these analysis techniques that unambiguously characterizes biological macromolecular slip, ideal, and catch-bond force-dependent unbinding. It deconvolutes the effects of how the experiment was performed with the unbinding force and time probably distributions, and it accounts for noise-based detection limits. We demonstrate the method’s ability to extract the molecular parameters from force-ramp optical trapping assay data, including the zero-force unbinding rate constant, force-dependent rate constants, and force sensitivities, using simulated data from various bond types.

## Theory

Consider a non-covalent, biological macromolecular bond, like one that underlies a protein-protein interaction. Based on classical thermodynamic models, the system has an unbinding constant, *k*_off_, at quasi-thermodynamic equilibrium (for bonds subject to loading that is slow compared to the timescales of force equilibrium and thermal fluctuations). In the absence of force and in a dilute solution, the average lifetime of a protein-protein interaction bond is the inverse of its unbinding constant, ⟨*τ*_off_⟩ = 1/*k*_off_. We consider an ensemble with *N* independent bonds formed at *t* = 0, of which *n*(*t*) of them remain bound at time *t*. The unbinding kinetics of the ensemble, assuming bonds do not reform after breaking, are

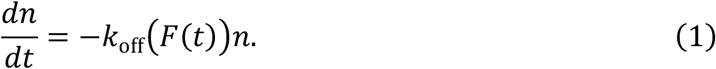

In general, the unbinding constant *k*_off_(*F*(*t*)) is a function of the force applied to the bond, and we consider the case that each bond is subject to a time-varying external force, *F*(*t*), e.g., as exerted by an optical tweezer.

By separating the variables and integrating Eq. (1), we find a cumulative distribution-like function for *n*(*t*),

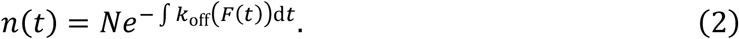

The functional form of *k*_off_(*F*(*t*)) depends on the bond type and how force changes with time, e.g., a linearly increasing force ramp (*F*(*t*) = *ft*) where *f*, the time rate of change of applied force, is constant. By applying models that relate the unbinding constant and force, we can complete the integration and obtain a functional form for the number of unbroken bonds as a function of time, *n*(*t*).

### Slip bonds

The unbinding constant of a slip bond, *k*_off,s_, increases exponentially with force per Bell’s model^16^

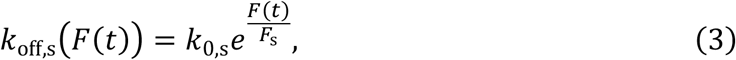

where *k*_0,s_ is the force-free unbinding constant and *F*_s_ is the force sensitivity of the slip bond. Slip bonds with small *F*_s_ as compared to the applied load are more force-sensitive than those with large *F*_s_. Thus, for a linearly increasing force ramp,

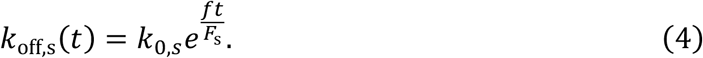

Therefore, Eq. (2) is

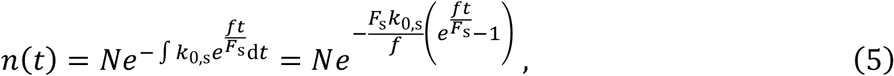

for a slip bond.

However, the fastest unbinding events may not be observable above experimental noise or may not be within the time sensitivity of force spectroscopy data. For example, binding/unbinding events in a linear force ramp experiment (Figure 1b) are only observable if the force at the time of dissociation is significantly larger than the amplitude of the thermal fluctuations of a bead in the trap times the effective stiffness of the trap, *F*_noise_. Therefore, we introduce a time scale, *t*_0_, representing the shortest observable event in a linear force ramp experiment, *t*_0_ = *F*_noise_/*f*. Thus, the total number of observable binding events is

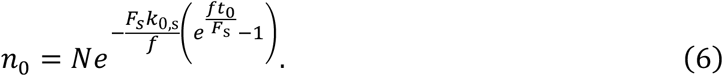

Due to the experimental noise, this analysis suggests that

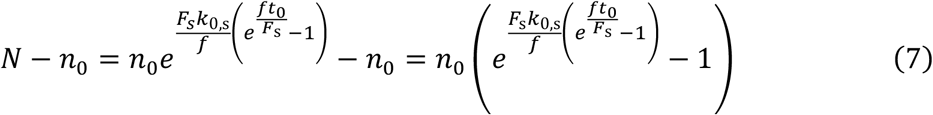

unbinding events occur in a time less than *t*_0_ and are missing from the observable dataset. Thus, to characterize the physical properties of a slip bond, one can record the number of bonds that remain unbroken as a function of time, and fit the data to

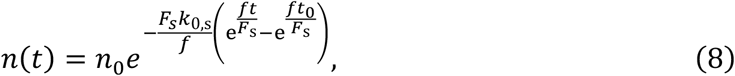

where the slip bond characteristics, *F*_s_ and *k*_0,*s*_, are the fitting parameters. *n*(*t*), and therefore *F*_s_ and *k*_0,s_, strongly depend on the loading rate, *f*, of the experiment. While *f* = *κv* can be controlled by changing with the stage translation rate, *v*, in a fixed-beam optical tweezer assay (Figure 1a) or trap translation in a fixed-stage optical tweezer assay, and the trap stiffness, *κ*_trap_, the loading rate is additionally complicated by the finite stiffness of the system, including linking molecules, *κ*_system_, where 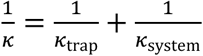.

### Ideal bonds

The unbinding constant of an ideal bond, *k*_off,i_, does not change with force. Hence, even when the applied force changes with time, the unbinding constant remains constant. In that case, the unbinding constant is *k*_off,i_(*t*) = *k*_0,i_ and Eq. (2) becomes

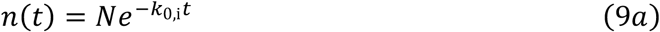

or

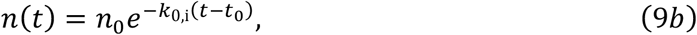

where *n*_0_ is the total number of observable binding events. *n*(*t*) is not a function of the loading rate, *f*, in the ideal bond case, and ideal bonds are special cases of slip bonds where the bond is entirely insensitive to force, i.e., for the limit when *F*_s_ → ∞ (Supplementary Information).

### Slip-ideal bonds

Recent data suggests that certain protein-protein interactions may be best modeled as a slip-ideal bond^28^. The force-dependent unbinding constant of a slip-ideal bond, *k*_off,s-i_(*F*(*t*)), increases exponentially with force per Bell’s model^16^ up to the slip-ideal transition force, *F*_s-i_, beyond which, it behaves as an ideal bond (Figure 2a). Therefore, the simplest mathematical model for the force-dependent unbinding constant of a slip-ideal bond is a piecewise function,

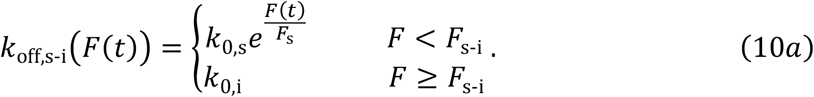

**Figure 2.**
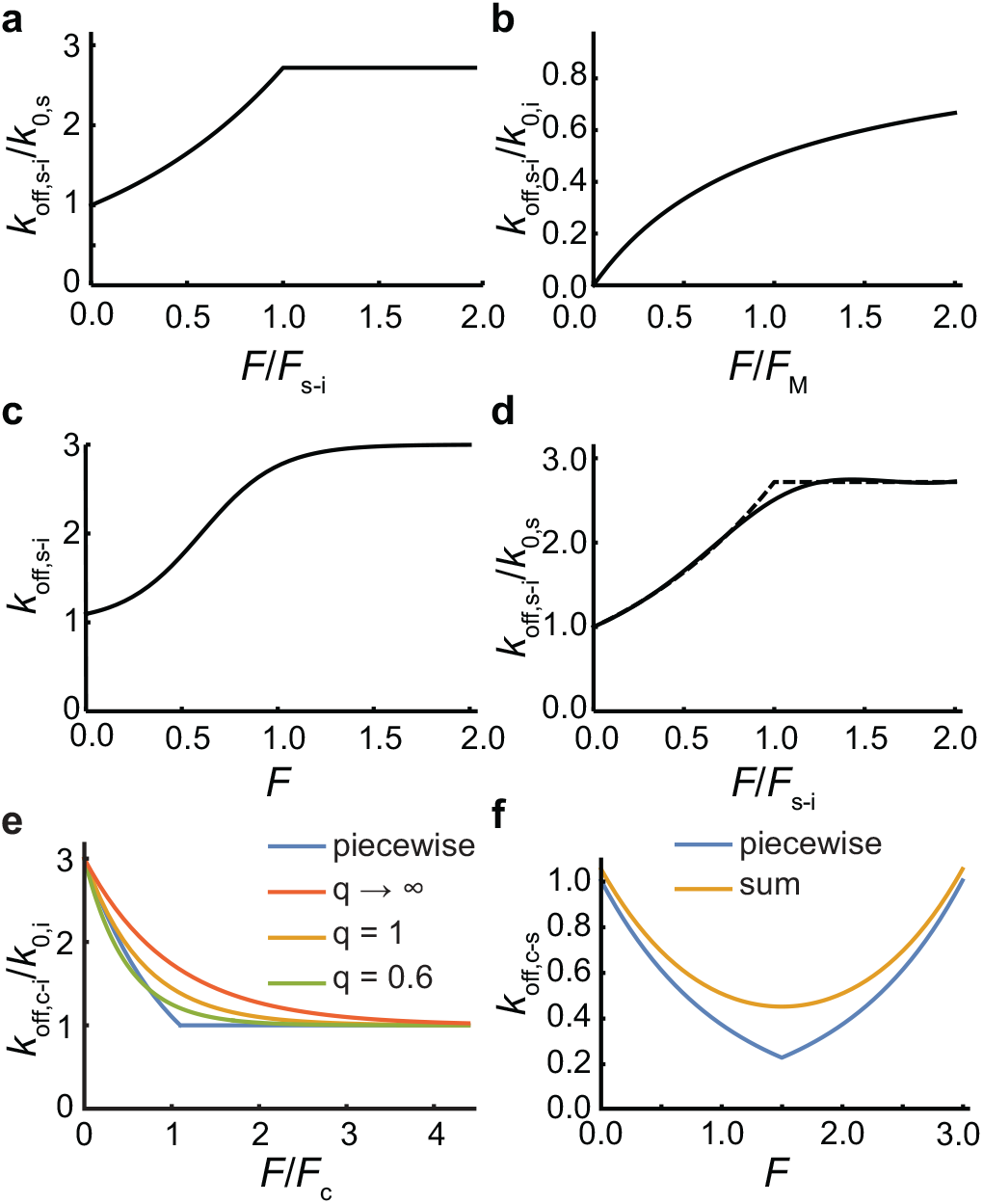
Analytical models for the force-dependent unbinding constant of slip-ideal, catch-ideal, and catch-slip bonds. **(a)** The piecewise function that explicitly models a slip-ideal bond (Eq. (10)) normalized by the unbinding constant at zero force and plotted as a function of force normalized by the slip-ideal transition force for the case of *F*_s-i_ = *F*_s_. **(b)** A Michaelis-Menten or Langmuir absorption-like approximation of the slip-ideal bond (Eq. (S1)) normalized by the ideal bond’s unbinding constant and plotted as a function of the force normalized by the Michaelis constant-like characteristic force. **(c)** A sigmoidal approximation of a slip-ideal bond (Eq. (S3)) where 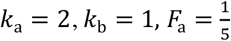 and 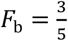. **(d)** A rational interpolation approximation (Eq. (S5), *solid line*) of the piecewise function from panel **(a)** (*dashed line*) for a slip-ideal bond with *F*_s-i_ = *F*_s_ = 1, which is equivalent to *β* = 1 (Supplementary information), normalized by the slip-bond unbinding constant, and plotted as a function of the force normalized by the characteristic force of the slip ideal transition, i.e., 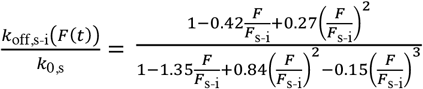 per Table S1. **(e)** The piecewise function (*blue*, Eq. (13)) that explicitly models a catch-ideal bond, as well as functional approximations of that function (*red, yellow, green*, Eq. (20)), normalized by the characteristic slip-bond unbinding constant and plotted as a function of force normalized by the force of the catch-bond for the representative case of *k*_0,c_ = 3*k*_0,i_. **(f)** The piecewise function (*blue*, Eq. (16)) and sum formulation (*yellow*, Eq. (17)) of a model for the force-dependent unbinding constant of the catch-slip bond. In both cases, we used *k*_0,c_ = 1 s^-1^, *F*_c_ = 1 pN, *k*_0,s_ = 0.05 s^-1^, and *F*_s_ = 1 pN as an example set of parameters. Given these parameters, *F*_c-s_ = 1.50 pN.

While there appear to be four independent parameters: the ideal bond’s unbinding constant, the slip bond’s unbinding constant and force sensitivity, and the slip-ideal transition force, *F*_s-i_, they are related by 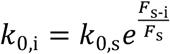, or equivalently *F*_s-i_ = *αF*_s_ where 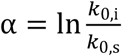, due to the continuity condition at *F* = *F*_s-i_. Thus, the slip-ideal model can be written as either

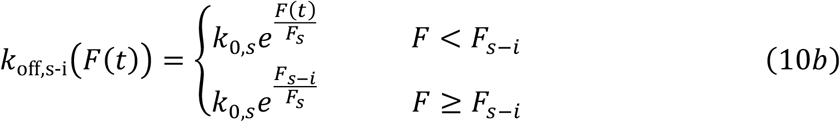

or

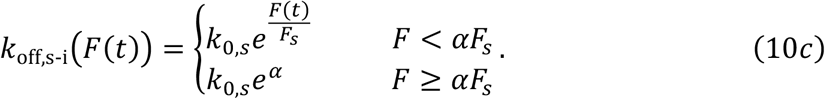

Unfortunately, the piecewise nature of Eq. (10) does not lend itself well to further mathematical analysis. However, there are multiple approaches to approximating slip-ideal bonds, which enable further analysis and ultimately fitting data to Eq. (2). These approaches include a Michaelis-Menten or Langmuir absorption-like function (Figure 2b)^31^, a sigmoidal model approximation (Figure 2c), and a rational interpolation^32,33^ approach (Figure 2c) that uses the RationalInterpolation function in Mathematica^34^, for example. See the Supporting Information for a more detailed discussion on these approximations of slip-ideal bonds.

### Catch bonds

The unbinding constant of the catch bond, *k*_off,c_, decreases exponentially with force per Bell’s model^16^,

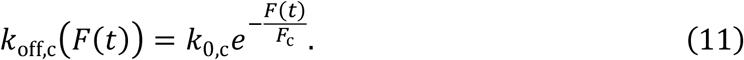

So, for a constant loading and *n*_0_ observable events, Eq. (2) becomes

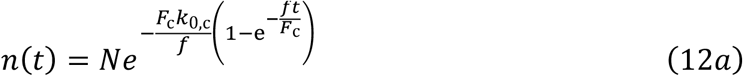

or

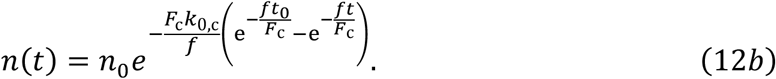

However, this is a relatively limited model because it models the unbinding rate constant as going to zero as *F* goes to infinity. Eq. (11) suggests that a catch bond will never break under an arbitrarily high load, which is non-physical. Therefore, Eq. (11) is only valid for low force. All known catch-bonds invariably transform into slip or ideal bonds as the forces increase beyond a critical value^35,36^. To capture the high-loading case, one can model all catch bonds as catch-slip or catch-ideal bonds.

### Catch-ideal bonds

A simple, theoretically appealing way to correct for the physical limitation of the catch bond is with a catch-ideal bond. A catch-ideal bond can be modeled as a bond that behaves like a catch bond under low loading and an ideal bond after the force crosses the catch-ideal transition force, *F*_c-i_. The unbinding constant of the catch-ideal bond, *k*_off,c-i_ (Figure 2e), is

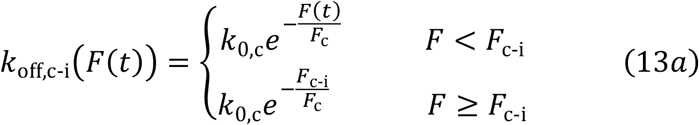

or

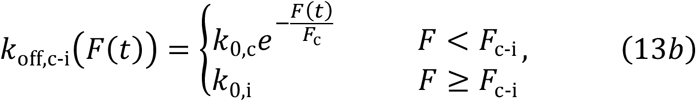

where only three of the four parameters are independent: 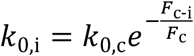 or 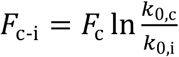 due to the continuity condition at *F* = *F*_c-i_.

We can approximate Eq. (13) with a sigmoidal function,

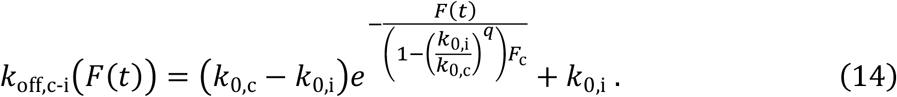

At zero force, the sigmoidal approximation behaves like a catch bond, *k*_off,c-i_(0) = *k*_0,c_, and *k*_off,c-i_(*F* → ∞) → *k*_0,i_ as force goes to infinity. Eq. (14) shares the same initial slope, 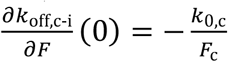, with Eq. (13) if *q* = 1, and it represents a simpler approximation, that 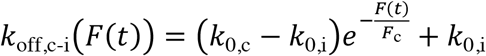, for q goes to infinity (Figure 2e). The best fit of Eq. (14) to the piecewise function (Eq. (13), found with a least-squares regression, Mathematica) occurs when *q* = 0.60 (Figure 2e). For constant loading rate, Eq. (2) becomes

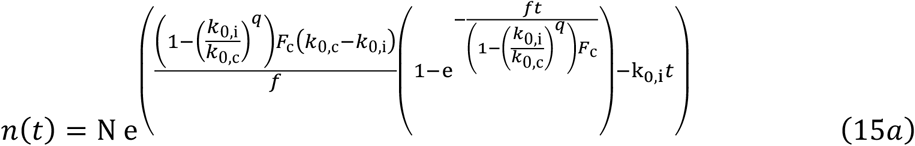

or

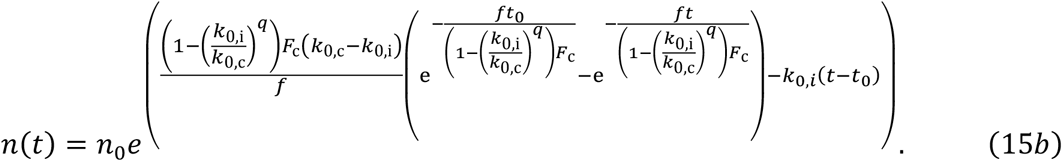

### Catch-slip bonds

While the catch-ideal bond is appealing, there is a stronger theoretical basis of^35,37^ and experimental evidence for^38^ catch-slip bonds. The conceptual model of a catch-slip bond suggests a transition from catch bond to slip bond behavior when the force exceeds a critical catch-slip transition value, *F*_c-s_. A five-parameter piecewise function can mathematically model catch-slip bonds (Figure 2f),

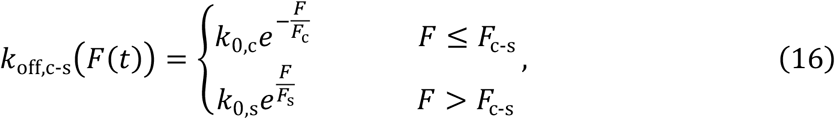

where only four parameters are independent because 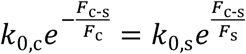, or equivalently 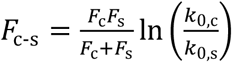, due to the continuity condition at *F* = *F*_c-s_.

Catch-slip bonds have been modeled as dissociating through one of two independent pathways: the catch bond pathway at relatively low force and short times, and the slip bond pathway at relatively high force and long times^10^. The analytical expression for this model is

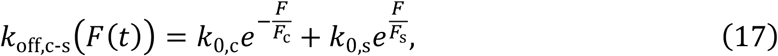

provided 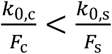 (Figure 2f). In this case, the catch-slip transition force, 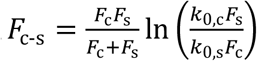, occurs when the first derivative of Eq. (17) with respect to force is zero. For constant loading rate, Eq. (2) becomes

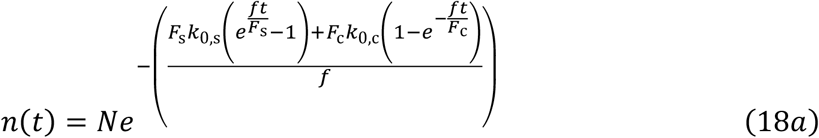

or

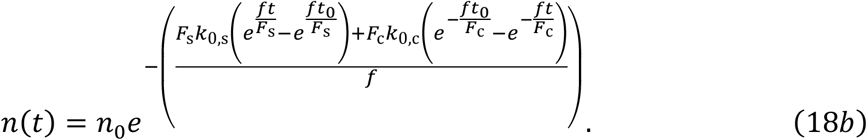

### Simulations

We stochastically simulated the unbinding of non-covalent, biomolecular bonds in MATLAB^39^ using the Gillespie algorithm^40^. We considered a system with *N*independently attached bonds at time *t* = 0 and applied a constantly increasing force (with constant loading rate *f*) to each bond. We simulated the time at which each bond, *i* = 1 → *N*, broke, *t*_*i*_ = *t*_*i*−1_ + *τ*_*i*_, where

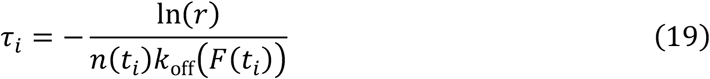

per standard implementation of the Gillespie algorithm^41^, *r* is a random number between 0 and 1 pulled from a uniform distribution, *n*(*t*_*i*_) = *N* − *i* is the number of bonds that remain attached after time *t*_*i*_, and *k*_off_(*F*(*t*_*i*_)) is the force-dependent unbinding constant.

## Results and discussion

We demonstrate the benefits of fitting single-molecule unbinding data to the cumulative distribution function-like *n*(*t*) over more traditional histogram analysis using simulated data. We do so, rather than using example experimental data, to demonstrate the power of this analysis on datasets for which we know the underlying biophysical nature of the bonds because we set them in the simulations. We use slip bonds and catch-slip bonds as examples (for brevity) because they are commonly reported bond types in the literature. However, the analysis could be easily extended to the other bond types and approximations of their functional forms, as discussed above and the Supporting Information.

### Slip bonds

We simulated *N* = 1000 slip bonds with unloaded unbinding rate *k*_0,s_ = 1 s^-1^ and force sensitivity *F*_1_ = 1 pN subjected to no loading (*k*_off,s_(*F* = 0) = *k*_0,s_ per Eq. (3) and 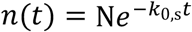, Figure 3a) and with linearly increasing loads (*k*_off,s_(*F* = *ft*) per Eq. (4) and *n*(*t*) per Eq. (5), Figure 3a). By comparing histograms of the time to detach for 1000 simulated bonds (Figure 3b), we found that the time to detachment distribution was a strong function of loading rate. With no external load (*F* = 0, Figure 3b, *upper left*), the histogram took the form of exponential decay, as expected for a single kinetic process. However, the characteristic form of the histogram became increasing Gaussian-like (though not strictly Gaussian) as the loading rate, *f*, increased (Figure 3b).

**Figure 3.**
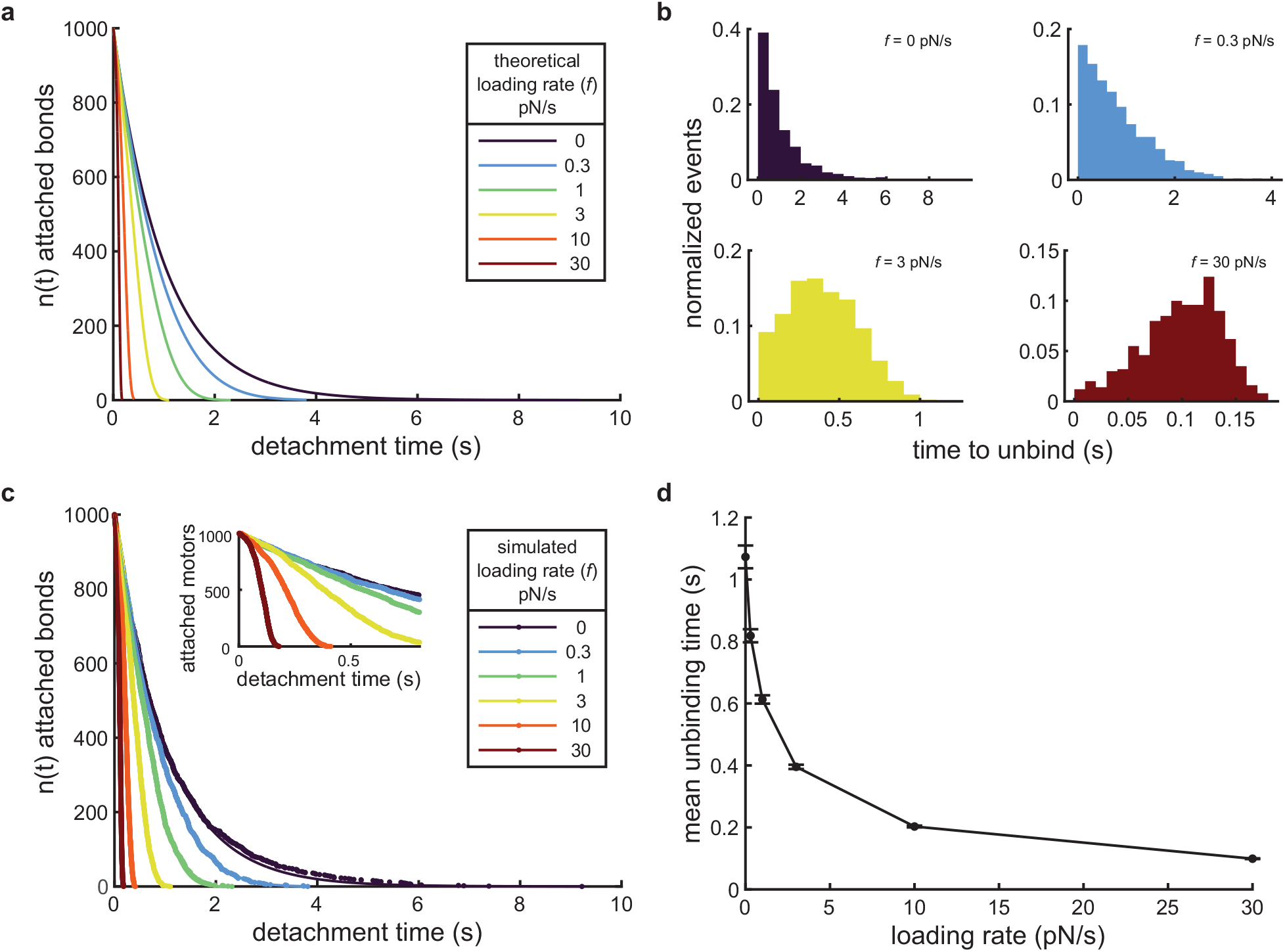
Calculating the force-dependent unbinding rate for slip bonds. **(a)** 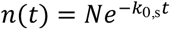 plotted for no loading (*dark purple line*) and Eq. (5) plotted for various loading rates (*colored lines*) with *N* = 1000 slip bonds that have an unloaded unbinding rate *k*_B,T_ = 1 s^-1^ and force sensitivity *F*_s_ = 1 pN. **(b)** Histograms of the time to unbind from a typical, *N* = 1000, slip bond simulation with parameters as in panel **(a)** for various loading rates, as indicated. **(c)** Number of bound slip bonds as a function of time from example simulations at various loading rates, as indicated. Inset shows detail highlighting the characteristic shape of *n*(*t*) for higher loading rates (3 pN/s *yellow*, 10 pN/s *orange*, and 30 pN/s *dark red*). The data (*dots*) were fitted to 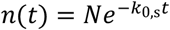 for *f* = 0 (*dark purple line*) and Eq. (5) for the others (*lines*). In both cases, see Table S2 for the fit parameters. The same example simulation data were used as in panel **(b)**, where applicable. **(d)** Mean unbinding time of 50 simulations, with *N* = 1000 slip bonds each, as a function of loading rate. The error bars represent s.e.m.

We also plotted the force-free and force-dependent dissociation of simulated slip bonds (same data as Figure 3b) as a function of their time to dissociate (*n*(*t*)). We fit these data to 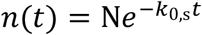 and Eq. (5) (Figure 3c, Table S2, and Supporting Information for details), respectively. We repeated the simulations 50 times and calculated the mean fitting parameters for each loading rate (Table 1). We found that the means of the fit parameters, *k*_0,*s*_ and *F*_0_, were not significantly different from the corresponding parameters used in the simulations (*p* values > 0.05 in all cases, Table 1, *t*-tests). Thus, our cumulative distribution function-like analysis uniquely and accurately determined the underlying biophysical parameters associated with slip bonds at each loading rate, suggesting one only needs to collect one such data set to characterize a slip bond.

**Table 1:** Mean fit parameters from 50 sets of *N* = 1000 simulated slip bond dissociation data sets.

| Loading rate (pN/s) | $k_{0,s}$ (s<sup>-1</sup>) (fit ± s.e.m.) | $p$-value | $F_s$ (pN) (fit ± s.e.m.) | $p$-value |
| --- | --- | --- | --- | --- |
| Simulation parameter | 1 | - | 1 | - |
| 0 | $1.009 \pm 0.005$ | 0.06 | - | - |
| 0.3 | $0.990 \pm 0.008$ | 0.21 | $1.019 \pm 0.038$ | 0.62 |
| 1 | $0.995 \pm 0.009$ | 0.58 | $0.998 \pm 0.015$ | 0.88 |
| 3 | $1.018 \pm 0.008$ | 0.06 | $1.010 \pm 0.008$ | 0.25 |
| 10 | $0.992 \pm 0.012$ | 0.48 | $0.997 \pm 0.006$ | 0.67 |
| 30 | $0.996 \pm 0.016$ | 0.81 | $1.002 \pm 0.006$ | 0.69 |

We also calculated the mean detachment time for each force-ramp loading rate condition (Figure 3d). In the unloaded case, the mean unbinding time was 1.073 s for the example data in Figure 3b, which was similar to but not a particularly great predictor of (7.3% error) the expected value of 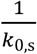 (1 s, for this simulation) for a first-order kinetic process (Supporting Information). However, when extending this analysis to the force-dependent unbinding properties, we found that the mean of the detachment time decreased with increasing loading rate (Figure 3d). Therefore, the underlying biophysical parameters of the bond are inaccessible to histogram analysis without explicitly accounting for the loading rate, and even then, it requires collecting large data sets at multiple loading rates (Supporting Information for more details).

### The effect of experimental noise on slip bonds

In an experiment (e.g., Figure 1a), it is impossible to distinguish single-molecule dissociation data for short times from the noise associated with the experiments. The noise “hides” detachment events if the detected data are smaller than four to five times the standard deviation of the noise. Selecting a detection threshold as high as five times the standard deviation is necessary to avoid misinterpreting the noise as a molecular unbinding event because the data collection rate necessary for high temporal resolution necessitates capturing many data points. For example, one would expect to misinterpret noise as a binding event 380 times with a 4σ threshold and 3.4 times with a 5σ threshold, on average, for 5 minutes of data collection at a 20 kHz sampling rate (6,000,000 data points). Depending on the total number of events within the measurement time, but frequently of order 10^2^-10^3^ events, hundreds of false-positive events could lead to a significant misinterpretation of the results. Given a loading rate, this detection threshold sets and effective minimum time for events, a detection threshold of *t*_0_, as described above.

We applied a detection threshold of *t*_0_ = 50 ms to the example simulated data in Figure 3b, and we found that the mean unbinding time in the unloaded case increased to 1.130 s from 1.073 s with *t*_0_ = 0 ms (*p* value = 0.28, *t*-test). However, in the case of *f* = 30 pN/s, we found that the mean unbinding time increased to 0.1075 s from 0.0991 s with *t*_0_ = 0 ms (*p* value < 0.0001, *t*-test). These results highlight the conclusion that the underlying biophysical parameters of the bond are in accessible in histogram analysis without specifically accounting for the loading rate and the detection limit (Supporting Information for more details).

We further probed the effect of the detection limit by discarding all data from our 50 simulations (same data as used in Table 1) with unbinding time *t* < *t*_0_ = 50 ms. We found that 50 to 110 data points of the *N* = 1000 were “hidden,” on average, by the detection limit (*N* − *n*_0_, Table 2), representing the shortest 5 to 11% of the events in the simulated data sets. The biased nature of the data loss (short events), and the strong effect of loading rate on the extent to which the data is lost (Table 2) makes estimating the number of “hidden” datapoints to from an experimental data set difficult. This data loss significantly affects any quantitative analysis of unbinding time histograms, as described above and detailed in the Supporting Information.

**Table 2.** **Mean fit parameters from the same 50 sets of simulated slip bond dissociation data as in Table 1, but where short events corresponding to those that are indistinguishable from experimental noise have been removed**

| Loading rate (pN/s) | $n_0$ (mean ± s.e.m.) | $k_{0,s}$ ($s^{-1}$) (fit ± s.e.m.) | $p$-value | $F_s$ (pN) (fit ± s.e.m.) | $p$-value |
| --- | --- | --- | --- | --- | --- |
| Simulation parameter | 1000 | 1 | - | 1 | - |
| 0 | $950.3 \pm 1.1$ | $1.008 \pm 0.005$ | 0.11 | - | - |
| 0.3 | $951.2 \pm 1.0$ | $0.990 \pm 0.009$ | 0.24 | $1.030 \pm 0.044$ | 0.49 |
| 1 | $950.6 \pm 1.0$ | $0.997 \pm 0.010$ | 0.79 | $1.003 \pm 0.016$ | 0.86 |
| 3 | $946.2 \pm 1.1$ | $1.013 \pm 0.010$ | 0.20 | $1.008 \pm 0.010$ | 0.41 |
| 10 | $937.1 \pm 1.1$ | $0.987 \pm 0.014$ | 0.34 | $0.995 \pm 0.007$ | 0.49 |
| 30 | $890.5 \pm 1.3$ | $0.984 \pm 0.022$ | 0.48 | $0.997 \pm 0.008$ | 0.74 |

However, we found that the situation was much improved when we fit these data to 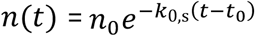 and Eq. (9) using the number of “detected” unbinding events in each simulation, *n*_0_, *f*, and *t*_0_ = 50 ms as fixed parameters. We calculated the mean of the 50 sets of fitting parameters for each loading rate (Table 2). We found that the means were not significantly different from the parameters used in the simulations, i.e., *k*_0,s_ = 1 s^-1^ and *F*_s_ = 1 pN (*p* values > 0.05 in all cases, Table 2, *t*-tests), and that none of the mean fitting parameters were significantly different from the fitting parameters found using all the data (*p* values > 0.05 in all cases, comparing data in Table 1 and Table 2, *t*-tests). Thus, despite the “hidden” data, we showed that our cumulative distribution function-like analysis unambiguously recovers the physical parameters of the slip bonds.

### Catch-slip bonds

We also simulated *N* = 1000 catch-slip bonds with an unloaded unbinding rate of *k*_0,c_ = 1 s^−1^, catch bond force sensitivity *F*_c_ = 1 pN, slip bond unloaded unbinding rate *k*_0,s_ = 0.05 s^−1^, and slip bond force sensitivity *F*_s_ = 1 pN (the same parameters used in Figure 2f, these equate to *F*_c-s_ = 1.5 pN) subjected to linearly increasing loads (*k*_off,c-s_(*F* = *ft*) per Eq. (16) and *n*(*t*) per Eq. (18), Figure 4a), as well as with no loading (*k*_off,c-s_(*F* = 0) = *k*_0,c_ per Eq. (16) and 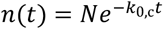, equivalent to the unloaded slip bond case, Figure 3a, *dark purple*). We found that the nature of the time to detach distribution histograms was a strong function of loading rate. Low external loading rates (*f* = 0.3 pN/s, Figure 4b, *upper left*) exhibited a nearly exponential distribution of unbinding times, like the other bond types we have discussed, and high external loading rates (*f* = 30 pN/s, Figure 4b, *lower right*) corresponded to a large Gaussian-like distribution of unbinding events at longer times. It is in the intermediate loading rate regime that two separate peaks, coming from the separate, significant contributions from the catch bond (short events) and slip bond (long events) behaviors, respectively, emerge (Figure 4b, *upper right* and *lower left*).

**Figure 4.**
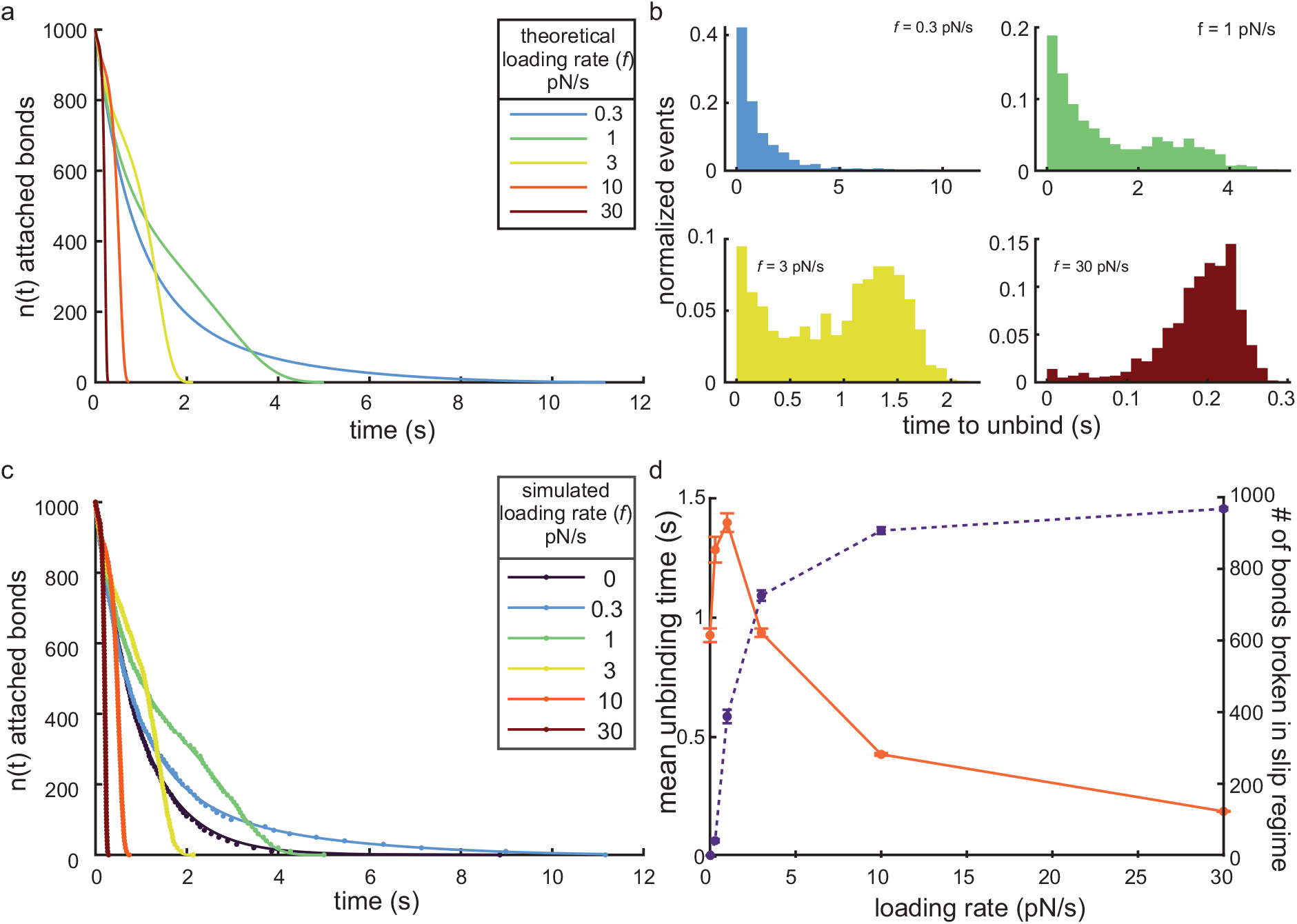
Calculating the force-dependent unbinding rate for catch-slip bonds. **(a)** Eq. (18a) plotted for various loading rates (colored lines) with *N* = 1000 slip bonds an unloaded unbinding rate of *k*_0,c_ = 1 s^A7^, catch bond force sensitivity of *F*_c_ = 1 pN, slip bond unbinding rate of *k*_0,s_ = 0.05 s^A7^, and slip bond force sensitivity of *F*_s_ = 1 pN, as in Figure 2f. **(b)** Histograms of the time to unbind from a typical *N* = 1000 slip bond simulation with parameters as in **(a)** for various loading rates, as indicated. **(c)** Number of bound catch-slip bonds as a function of time for example simulations at various loading rates, as indicated. The data (*dots*) were fit to 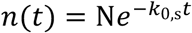 for *f* = 0 (*dark purple line*) and Eq. (18a) for the others (*lines*). The equations were well-fit to the data in all of the example cases shown here, see Table S3 for the fit parameters and Supporting Information for more detail. The same example simulation data were used as in panel **(b)**, where applicable. **(d)** Mean unbinding time (*orange solid line*, left axis) and mean number of bonds broken in the slip bond regime (*purple dashed line*, right axis) of 50 simulations, with *N* = 1000 catch-slip bonds each, as a function of loading rate. The error bars represent the s.e.m. on both data sets.

We also plotted the force-free and force-dependent dissociation of simulated slip bonds (same data as Figure 4b) as a function of their time to dissociate (*n*(*t*)). We fit these data to 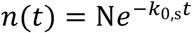 and Eq. (18a) (Figure 4c, Table S3, and Supporting Information for details), respectively. As we did with slip bonds, we input the values for *N*and *f* as fixed parameters in the fit and repeated the simulations 50 times. Unlike the case of slip bonds, we found that non-linear least squares regression was unable to uniquely determine fitting parameters in multiple of the simulated data sets at lower and higher loading rates (Supporting Information). Of the 50 simulations, we found that only 22 of the simulated data sets were well-fit by Eq. (18a) for low loading rate (0.3 pN/s) and 40 of the simulated data sets were well-fit by Eq. (18a) for high loading rate (30 pN/s).

We suspected that the errors in the fits were due to an “overfitting” of the data. To investigate this hypothesis, we determined whether the force at unbinding was above for below the catch-slip transition force, *F*_c-s_, for each unbinding event to classify whether the bond dissociated in the catch (*F* < *F*_c-s_) or slip (*F* > *F*_c-s_) regime. We plotted the mean number of bonds dissociating in the slip regime for the 50 simulations with *N* = 1000 total bonds (Figure 4d, *purple dashed line*). We found that most of the bonds dissociated in the catch bond regime (Figure 4d, *purple dashed line*) when the loading rate was low, corresponding to an exponential-like distribution of unbinding events (Figure 4b, upper left). At high loading rates, the fraction of bonds that dissociated in the slip force regime was high (Figure 4d, *purple dashed line*), corresponding to a large Gaussian-like distribution at relatively longer times (Figure 4b, lower right) and similar to slip bonds (Figure 3b, lower right). Together, these data suggest that the slip bond parameters have little effect on the unbinding behaviors at low loading rates and that the catch bond parameter have little effect on the unbind behaviors at high loading rates. Thus, attempting to fit Eq. (18a), which contains both the slip and catch bond-related parameters, is less effective at low or high loading rates.

Moreover, even when the low and high loading rate data are well-fit by Eq. (18a), i.e., the non-linear least squares regression was able to uniquely determine fitting parameters, we found that the standard errors of those fits tend to be significantly larger than for intermediate loading rates (Tables 3 and S3, and Supporting Information for more details). Therefore, these results suggest that one must use intermediate loading rates, i.e., loading rates for which a significant number of bonds dissociate in the catch and slip regimes (Figure 4d, *purple dashed line*), to determine the biophysical properties of catch-slip bonds.

**Table 3.** Mean fit parameters from 50 sets of *N* = 1000simulated catch-slip bond dissociation data sets.

| Loading rate (pN/s) | Significant fits | $k_{0,c}$ ($s^{-1}$)<br>(mean ± s.e.m.) | $p$ value | $F_c$ (pN)<br>(mean ± s.e.m.) | $p$ value | $k_{0,s}$ ($s^{-1}$)<br>(mean ± s.e.m.) | $p$ value | $F_s$ (pN)<br>(mean ± s.e.m.) | $p$ value |
| --- | --- | --- | --- | --- | --- | --- | --- | --- | --- |
| Simulation parameter | 50 | 1 | - | 1 | - | 0.05 |  | 1 | - |
| 0 | 50 | $1.0478^* \pm 0.0054$ | 0.69 | - | - | - | - | - | - |
| 0.3 | 22 | $1.0058 \pm 0.0171$ | 0.74 | $1.1698 \pm 0.0698$ | 0.02 | $0.0360 \pm 0.0160$ | 0.39 | $0.6043 \pm 0.1441$ | 0.01 |
| 1 | 50 | $1.0115 \pm 0.0109$ | 0.30 | $0.9826 \pm 0.0342$ | 0.61 | $0.0589 \pm 0.0048$ | 0.07 | $1.0341 \pm 0.0304$ | 0.27 |
| 3 | 50 | $1.0146 \pm 0.0145$ | 0.32 | $1.0467 \pm 0.0295$ | 0.12 | $0.0485 \pm 0.0017$ | 0.38 | $0.9878 \pm 0.0093$ | 0.20 |
| 10 | 47 | $1.0965 \pm 0.0326$ | 0.005 | $1.0120 \pm 0.0547$ | 0.82 | $0.0496 \pm 0.0016$ | 0.98 | $0.9952 \pm 0.0065$ | 0.46 |
| 30 | 40 | $1.3684 \pm 0.0795$ | < 0.0001 | $0.8083 \pm 0.0924$ | .04 | $0.0502 \pm 0.0021$ | 0.92 | $0.9974 \pm 0.0070$ | 0.71 |

\* $k_{off,c-s}(F = 0) = k_{0,c} + k_{0,s} = 1.05$ per Eq. (17) for the parameters used in the simulation

We also calculated the mean detachment time for each force-ramp loading rate condition (Figure 4d, *orange solid line*). We found that the mean detachment time increased for faster loading rates while the dissociations were dominated by the catch-bond behavior (Figure 4d, *purple dashed line*). However, this trend reversed at an intermediate loading rate and the mean detachment time decreased with increasing loading rate (Figure 4d, *orange solid line*) as more the dissociations were dominated by the slip-bond behavior (Figure 4d, *purple dashed line*). We had found that determining the underlying biophysical parameters of slip bonds from characterizations of the mean unbinding time and associated histograms requires collecting large data sets at multiple loading rates (Supporting Information for more details). However, we found that, to the best of our knowledge, an equivalent analysis did not yield analytical solutions, even with special functions, from which we could calculate the underlying biophysical parameters of catch-slip bonds using the mean unbinding time and associated histograms.

Together, this analysis and these data highlight the importance of performing single-molecule experiments at optimized experimental conditions, i.e., ones that enable characterization of all the physics associated with catch-slip bonds, and the utility of our cumulative distribution-like analysis.

### Effect of experimental noise on catch-slip bonds

As discussed above, noise in force-spectroscopy data can hide dissociations that occur at short times. To investigate the role of experimental noise on the analysis of catch-slip bonds, we excluded simulated dissociation events from our 50 simulations (same data as used in Table 3) with unbinding time *t* < *t*_0_ = 50 ms. We fit *n*(*t*) with Eq. (18b) to calculate the corresponding bond parameters (Table 4). We found that non-linear least squares regression uniquely determined fitting parameters in fewer of the simulated data sets, particularly at higher loading rates, than when not accounting for experimental noise (Supporting Information). Of the 50 simulations, we found that only 24 of the *f* = 0.3 pN/s, 33 of the *f* = 10 pN/s, and 5 of the *f* = 30 pN/s simulated data sets were well-fit by Eq. (18b) (Table 4).

**Table 4.** **Mean fit parameters from the same 50 sets of simulated catch-slip bond dissociation data as in Table 3, but where short events corresponding to those that are indistinguishable from experimental noise have been removed**

| Loading rate (pN/s) | Significant fits | $n_0$<br>(mean ± s.e.m.) | $k_{0,c}$ (s<sup>-1</sup>)<br>(mean ± s.e.m.) | $p$<br>value | $F_c$ (pN)<br>(mean ± s.e.m.) | $p$<br>value | $k_{0,s}$ (s<sup>-1</sup>)<br>(mean ± s.e.m.) | $p$<br>value | $F_s$ (pN)<br>(mean ± s.e.m.) | $p$<br>value |
| --- | --- | --- | --- | --- | --- | --- | --- | --- | --- | --- |
| Simulation parameter | 50 |  | 1 | - | 1 | - | 0.05 | - | 1 | - |
| 0 | 50 | 949.4 ± 1.1 | 1.0484* ± 0.0053 | 0.76 | - | - | - | - | - | - |
| 0.3 | 24 | 948.4 ± 0.8 | 0.9956 ± 0.0164 | 0.79 | 1.1764 ± 0.0678 | 0.02 | 0.0391 ± 0.0150 | 0.47 | 0.6217 ± 0.1397 | 0.01 |
| 1 | 50 | 949.3 ± 1.0 | 1.0149 ± 0.0128 | 0.25 | 0.9944 ± 0.0416 | 0.89 | 0.0594 ± 0.0050 | 0.06 | 1.0348 ± 0.0317 | 0.28 |
| 3 | 50 | 950.6 ± 0.7 | 0.9972 ± 0.0214 | 0.90 | 1.1124 ± 0.0449 | 0.02 | 0.0472 ± 0.0018 | 0.13 | 0.9812 ± 0.0095 | 0.05 |
| 10 | 33 | 957.0 ± 0.9 | 1.3098 ± 0.1197 | 0.01 | 1.1293 ± 0.1145 | 0.27 | 0.0482 ± 0.0017 | 0.30 | 0.9897 ± 0.0075 | 0.18 |
| 30 | 5 | 967.7 ± 0.6 | 0.9693 ± 0.3377 | 0.93 | 0.7971 ± 0.2214 | 0.41 | 0.0541 ± 0.0049 | 0.45 | 1.0131 ± 0.0159 | 0.46 |

\* $k_{\text{off},c-s}(F = 0) = k_{0,c} + k_{0,s} = 1.05$ per Eq. (17) for the parameters used in the simulation

As we observed for the simulated data without accounting for the effect of experimental noise (Table 3), the cumulative distribution function-like analysis technique was able to resolve the catch-bond fitting parameters reasonably well (despite a 17%, *p* value = 0.02, error in resolving the force sensitivity of the catch bond) with low s.e.m. of the fits in low loading rate simulations, and the slip-bond fitting parameters (despite a low number of well-fit data sets) in the high loading rate simulation (Table 4). Additionally, as we observed for the simulated data without accounting for experimental noise (Table 3), catch-bond fitting parameters were not well resolved at high loading rate simulations, and slip-bond fitting parameters were not well resolved at low loading rates (Table 4). Like we found with slip bonds, none of the fit parameters differed significantly from those found when accounting for the complete simulated data set (p values > 0.05, two-sample *t*-tests, Tables 3 and 4), again highlighting the benefit of our method when analyzing noisy experimental data.

These data again serve to highlight the effectiveness of the cumulative distribution-like analysis to resolve the biophysical parameters associated with catch-slip bonds, even when experimental noise hides short events. However, the reduction in the number of well-fit data sets, and the increased sensitivity of the catch bond parameters to the loading rate when short events are hidden by noise, does strongly suggest that loading rates must be chosen carefully, particularly for catch-slip bonds with relatively low catch-slip transition force, *F*_c-s_.

## Conclusion

We presented an analytically derived cumulative distribution-like function of unbinding events (*n*(*t*)) for various biological macromolecular bond types when subject to force. We showed how an *n*(*t*), cumulative distribution function-like, based approach can be used to analyze force-dependent dissociation force spectroscopy data. We demonstrated the benefits and limitations of the technique using stochastic simulations (Gillespie algorithm) of slip and catch-slip bonds. The approach can determine the detachment rate and force sensitivity of biological macromolecular bonds from force spectroscopy experiments by explicitly accounting for loading rate more efficiently than histogram-based analyses. This analysis approach requires fewer, smaller data sets than alternative approaches. Additionally, the approach returns similar (not statistically different) results when short events are hidden by noisy data. We suggest that this approach provides an improved systematic and quantitative method to distinguishing various bond types and characterizing their underlying biophysical properties.

We also analyzed the effect of using a range of loading rates to probe the force-dependent unbinding of biological macromolecules. Our simulated data suggests that this analysis of slip-bonds is largely insensitive to loading rate. However, care must be taken to ensure a significant fraction of the bonds dissociate in both the catch- and slip-bond force regimes, when the bond is a catch-slip bond. Thus, if an experiment is being done on a bond of unknown type, multiple loading rates are necessary to ensure all possible molecular dissociation pathways are sufficiently sampled in the experiments to resolve their underlying biophysical parameters.

In summary, our approach provides a framework for an improved analysis of force-dependent biological macromolecular dissociation force spectroscopy data. It explicitly accounts for the loading rate, which may be complicated by optical tweezer trap stiffness and biological macromolecular stiffness, to distinguish between and fully characterize the biophysical properties of protein-protein and protein-ligand bonds.

## Acknowledgements

AP was supported by a Clemson Research Fellows R-Initiative Award from Clemson University. This work was supported by the National Institute of Allergy and Infectious Diseases (NIAID) of the National Institutes of Health under award number R15AI137979, by the National Institute of General Medical Sciences (NIGMS) of the National Institutes of Health under award number P30 GM131959, and by Clemson University. We would like to thank Ashok Pabbathi for his sample data (Figure 1b) and Ashok Pabbathi and Subash Godar for their insightful discussions. Additionally, we thank Marija Zanic for reading and commenting on early drafts of the manuscript.

## Author contributions

AP contributed to the conceptualization of the study, performed the analytical and computational work, analyzed the results, and drafted the manuscript. JA contributed to the conceptualization of the study, performed the analytical and computational work, analyzed the results, revised the manuscript, and supervised the project.

