## Supplementary Information for "Calculating the force-dependent unbinding rate of biological macromolecular bonds from the force-ramp optical trapping assays"

#### Supplementary Theory

##### Ideal bonds are special cases of slip bonds where the bond is entirely insensitive to force

The unbinding constant of a slip bond, $k_{\text{off,s}}$, increases exponentially with force per Bell's model[1] (Eq. 3, main text). When we take the limit of the slip bond being entirely insensitive to force, i.e., for the limit when $F_{\text{s}} \to \infty$, the exponent of the exponential term in Eq. 3 approaches 0 and thus the exponential term approaches 1. Therefore, in the entirely insensitive limit $\lim_{F_s \to \infty} k_{\text{off,s}}(F(t)) = k_{0,\text{s}}$ takes the functional form of an ideal bond ($k_{\text{off,i}}(t) = k_{0,\text{i}}$). So, ideal bonds are just special cases of slip bonds that are entirely insensitive to force.

##### Slip-ideal bond approximations

Berger et al.[2] approximated the slip-ideal bond as a Michaelis-Menten or Langmuir absorption-like function

$$k_{\text{off,s-i}}(F(t)) = \frac{k_{0,\text{i}} F}{F_{\text{M}} + F} \qquad (S1a)$$

or

$$k_{\text{off,s-i}}(F(t)) = \frac{k_{0,\text{i}} \frac{F}{F_{\text{M}}}}{1 + \frac{F}{F_{\text{M}}}} \qquad (S1b)$$

where the Michaelis constant-like $F_{\text{M}}$ accounts for both the slip-ideal transition force and the force sensitivity of the slip bond. Smaller values of $F_{\text{M}}$ simultaneously correspond to a lower slip-ideal transition force and higher force sensitivity of the slip bond than larger values of $F_{\text{M}}$. Based on the Michaelis-Menten-like approximation of the slip-ideal bond and assuming a linear force ramp, Eq. 2 from the main text becomes

$$n(t) = N e^{-k_{0,\text{i}} t} \left( \frac{F_{\text{M}}}{F_{\text{M}} + f t} \right)^{-\frac{F_{\text{M}} k_{0,\text{i}}}{f}}. \qquad (S2a)$$

or

$$n(t) = n_0 e^{-k_{0,\mathrm{i}}(t-t_0)} \left(\frac{F_\mathrm{M} + f t_0}{F_\mathrm{M} + f t}\right)^{-\frac{F_\mathrm{M} k_{0,\mathrm{i}}}{f}} . \qquad (S2b)$$

where $n_0$ is the total number of observable binding events.

While the Michaelis-Menten-like model approximates the data reported by Nicholas et al.[3], it lacks some fundamental characteristics of the slip-ideal bond (Eq. 10 from the main text), i.e., the unbinding constant, $k_{\mathrm{off,s\text{-}i}}$, is 0 at $F = 0$ rather than $k_{0,\mathrm{s}}$, and the second derivative of $k_{\mathrm{off,s\text{-}i}}(F(t))$ with force is negative rather than positive at forces below the slip-ideal transition force. The former could be accounted for easily by adding a constant to Eq. S1, but the latter is a fundamental feature of the Michaelis-Menten-like approximation. These represent significant deviations from the piecewise slip-ideal function, which is rooted in Bell's model.

An alternative approximation is to use a sigmoidal approximation

$$k_{\mathrm{off,s\text{-}i}}(F(t)) = \frac{k_\mathrm{a}}{1 + e^{-\frac{(F - F_\mathrm{b})}{F_\mathrm{a}}}} + k_\mathrm{b} \qquad (S3)$$

where $F_\mathrm{a}$ relates to the force sensitivity of the slip bond, $F_\mathrm{b}$ relates to the slip-ideal transition force, $k_\mathrm{a}$ most closely relates to $k_{0,\mathrm{i}}$, and $k_\mathrm{b}$ most closely relate to $k_{0,\mathrm{s}}$. However, the details of these relationships are convoluted with each other ($k_\mathrm{a} = k_{0,\mathrm{i}} - k_\mathrm{b}$ and $k_\mathrm{b} = \frac{k_{0,\mathrm{s}} - \frac{k_{0,\mathrm{i}}}{1+e^{-\frac{F_\mathrm{b}}{F_\mathrm{a}}}}}{1 - \frac{1}{1+e^{-\frac{F_\mathrm{b}}{F_\mathrm{a}}}}}$). Based on the sigmoidal approximation of the slip-ideal bond and assuming a linear force ramp, Eq. 2 from the main text becomes

$$n(t) = N e^{-k_\mathrm{b} t} \left[\frac{1 + e^{\frac{F_\mathrm{b}}{F_\mathrm{a}}}}{e^{\frac{F_\mathrm{b}}{F_\mathrm{a}}} + e^{\frac{ft}{F_\mathrm{a}}}}\right]^{\frac{F_\mathrm{a} k_\mathrm{a}}{f}} \qquad (S4a)$$

and

$$n(t) = n_0 e^{-k_\mathrm{b}(t-t_0)} \left[\frac{e^{\frac{F_\mathrm{b}}{F_\mathrm{a}}} + e^{\frac{f t_0}{F_\mathrm{a}}}}{e^{\frac{F_\mathrm{b}}{F_\mathrm{a}}} + e^{\frac{ft}{F_\mathrm{a}}}}\right]^{\frac{F_\mathrm{a} k_\mathrm{a}}{f}} . \qquad (S4b)$$

Another approach to approximating the slip-ideal bond (Eq. 10 from the main text) is to perform rational interpolation[4,5] using the RationalInterpolation function in Mathematica[6], for example. Rational interpolation of Eq. 10 from the main text with the numerator of order $m = 2$ and denominator of order $n = 3$ over the interval of external force from 0 to twice the slip-ideal transition force ($2F_{\mathrm{s\text{-}i}}$) yields an expression of the form

$$k_{\mathrm{off,s\text{-}i}}(F(t)) = k_{0,s} \frac{c_0 + c_1 \bar{F} + c_2 \bar{F}^2}{c_3 + c_4 \bar{F} + c_5 \bar{F}^2 + c_6 \bar{F}^3} \qquad (S5)$$

where $\bar{F} = \frac{F}{F_{\text{s-i}}}$ and the fit coefficients are a function of $\frac{F_{s-i}}{F_s} = \ln\frac{k_{0,\text{i}}}{k_{0,\text{s}}} = \beta$ (Table S1). Based on the rational interpolation approximation of the slip-ideal bond and assuming a linear force ramp, Eq. 2 must be solved numerically.

**Table S1. Constants ($c_i$s) from the rational interpolation of the force-dependent slip-ideal unbinding constant**

| $\beta$ | $c_0$ | $c_1$ | $c_2$ | $c_3$ | $c_4$ | $c_5$ | $c_6$ |
|---|---|---|---|---|---|---|---|
| 0.25 | 1 | -3.32 | 6.68 | 1 | -4.25 | 9.33 | -3.35 |
| 0.5 | 1 | -1.41 | 1.46 | 1 | -2.34 | 2.72 | -0.74 |
| 1 | 1 | -0.42 | 0.27 | 1 | -1.35 | 0.84 | -0.15 |
| 2 | 1 | 0.14 | 0.07 | 1 | -0.80 | 0.26 | -0.26 |
| 4 | 1 | 0.42 | 0.24 | 1 | -0.46 | 0.076 | -0.0036 |

While the rational interpolation approach to approximating the slip-ideal bond is appealing, it does not lend itself to the cumulative distribution function-like analysis of single-molecule force-dependent dissociation experiments as the constants (Table S1) are highly dependent on $\beta = \ln\frac{k_{0,\text{i}}}{k_{0,\text{s}}}$, the approximation cannot be used outside the interval of normalized force $\bar{F} = \frac{F}{F_{\text{s-i}}}$ for which it was interpolated ($0 < \bar{F} < 2$ in the case of Table S1), and Eq. 2b must be integrated numerically.

### Supplementary results

#### Slip bond simulations

**Table S2. Fit parameters from example simulated slip bond dissociation data, shown in Figure 3c.**

| Loading rate (pN/s) | $k_{0,\text{s}}$ ($\text{s}^{-1}$) (fit ± s.e. of the fit) | $F_\text{s}$ (pN) (fit ± s.e. of the fit) |
|---|---|---|
| Simulation parameter | 1 | 1 |
| 0 | 0.9797 ± 0.0001 | - |
| 0.3 | 0.9363 ± 0.0001 | 0.8308 ± 0.0043 |
| 1 | 0.9014 ± 0.0018 | 0.8677 ± 0.0037 |
| 3 | 0.8884 ± 0.0025 | 0.9110 ± 0.0027 |
| 10 | 0.9718 ± 0.0022 | 0.9974 ± 0.0014 |
| 30 | 1.0586 ± 0.0038 | 1.0349 ± 0.0015 |

We determined the fit parameters for simulated slip bond dissociation data (Figure 3 and Tables 1 and S2) with the fit (nonlinear least squares method) function in MATLAB. The values represent the coefficients ± the standard error of those fits to $n(t) = \text{N}e^{-k_{0,\text{s}}t}$ for $f = 0$ and Eq. (5) in the main text. We input the values for $N = 1000$ and the loading rate $f$ as fixed parameters in the fit rather than free fitting parameters because the total number of unbinding events detected and the loading rate are known with reasonable confidence in a force spectroscopy experiment, i.e., optical tweezers. In each example case (Table S2), the fit parameters were significantly different from the value used to stochastically generate the simulated data, $k_{0,\text{s}} =$

1 $s^{-1}$ and $F_s = 1$ pN, *p* values < 0.0001 in all cases (smaller than the double precision data type in MATLAB).

We also determined the fit parameters for simulated slip bond dissociation data with short events, corresponding to those that are indistinguishable from the experimental noise, removed (Table 2) using the fit (nonlinear least squares method) function in MATLAB to $n(t) = n_0 e^{-k_{0,s}(t-t_0)}$ and Eq. (8) in the main text. Again, we used the number of "detected" unbinding events in each simulation, $n_0$, $f$, and $t_0 = 50$ ms as fixed parameters in the fitting because the total number of unbinding events detected that were distinguishable from the noise, $n_0$, and the loading rate, $f$, are known with reasonable confidence in a force spectroscopy experiment, i.e., optical tweezers.

#### Expected value for average slip bond detachment time using histogram analysis

We calculated the expected value, *E(f)*, of the time to dissociate for slip bond detachment as a function of loading rate,

$$E(f) = \int_0^\infty t\,p(t)\,dt \qquad (S6)$$

where $p(t)$ is the probability distribution function of dissociation time. $p(t)$ is the first derivative of the cumulative distribution function dissociation times $p(t) = \frac{1}{N}\frac{d}{dt}\{\mathrm{cdf}(t)\}$ and $\mathrm{cdf}(t) = N - n(t)$. Thus, to account for the loading rate in the histogram analysis, we found that the mean of the detachment time of a slip bond as a function of loading rate is

$$E(0) = \frac{1}{k_{0,s}} \qquad (S7)$$

for the unloaded case (as expected) and

$$E(f) = \frac{F_s}{f} e^{\frac{F_s k_{0,s}}{f}} \Gamma\left(0, \frac{F_s k_{0,s}}{f}\right) \qquad (S8)$$

for the loaded case, where $\Gamma(0, \mathrm{x})$ is the upper incomplete gamma function of $x$. While this analysis is possible, and the loading rate correction can be applied, because gamma functions do not provide much insight to most people, we find that fitting the data to $n(t) = \mathrm{N}e^{-k_{0,s}t}$ and Eq. 5 is a more intuitive procedure. Additionally, this analysis requires collection of data at multiple rates to deconvolve loading rate from the results because Eq. S8 is a single equation with two unknowns, $k_{0,s}$ and $F_s$. So, while directly fitting to $n(t) = \mathrm{N}e^{-k_{0,s}t}$ and Eq. 5 can yield $k_{0,s}$ and $F_s$ with a single, well designed, force spectroscopy experiment, taking a histogram of unbinding time (or force) approach requires at least two independent loading rates to deconvolve the effect of the experimental procedures from the results.

In principle, the need for collecting data at two independent loading rates could be mitigated by fitting the histogram data to $p(t)$, directly. Thus, one would fit the histogram to

$$p(t) = k_{0,s} e^{-k_{0,s}t}, \qquad (S9)$$

for the unloaded case, and

$$p(t) = k_{0,s} e^{-\frac{F_s k_{0,s}}{f}\left(e^{\frac{ft}{F_s}}-1\right)+\frac{ft}{F_s}} \qquad (S10)$$

for a constant loading rate $f$. However, this requires fitting to a histogram of data which reduces the degrees of freedom, and thus the confidence of the fit, due to binning of the data. So, to get the same level of confidence in the fit parameters, $k_{0,s}$ and $F_s$, one would have to collect an order of magnitude more data.

The situation is further complicated in the case of "missing events," when experimental noise hides the shortest binding events. In this case, we found that the mean of the detachment time of a slip bond as a function of loading rate is

$$E(0) = \frac{1 + k_{0,s} t_0}{k_{0,s}} \qquad (S11)$$

for the unloaded case and

$$E(f) = t_0 - \frac{F_s}{f} e^{\frac{F_s k_{0,s} e^{\frac{f t_0}{F_s}}}{f}} \mathrm{Ei}\left(-\frac{F_s k_{0,s} e^{\frac{f t_0}{F_s}}}{f}\right) \qquad (S12)$$

for the loaded case where $\mathrm{Ei}(x)$ is the exponential integral of $x$.

#### Catch-slip bond simulations

**Table S3. Fit parameters from example simulated catch-slip bond dissociation data, shown in Figure 4c.**

| Loading rate (pN/s) | $k_{0,c}$ ($s^{-1}$) (fit ± s.e. of the fit) | $p$ value | $F_c$ (pN) (fit ± s.e. of the fit) | $p$ value | $k_{0,s}$ ($s^{-1}$) (fit ± s.e. of the fit) | $p$ value | $F_s$ (pN) (fit ± s.e. of the fit) | $p$ value |
|---|---|---|---|---|---|---|---|---|
| Simulation parameter | 1 | | 1 | | 0.05 | | 1 | |
| 0 | 1.0576 ± 8.0840 x $10^{-4}$ | < 0.0001 | - | - | - | - | - | - |
| 0.3 | 1.0537 ± 0.0306 | 0.08 | 0.8268 ± 0.0406 | < 0.0001 | 0.0735 ± 0.0318 | 0.46 | 1.4377 ± 0.4585 | 0.34 |
| 1 | 0.9909 ± 0.0014 | < 0.0001 | 1.1726 ± 0.0095 | < 0.0001 | 0.0222 ± 0.0008 | < 0.0001 | 0.7732 ± 0.0074 | < 0.0001 |
| 3 | 1.0784 ± 0.0046 | < 0.0001 | 1.1196 ± 0.0122 | < 0.0001 | 0.0323 ± 0.0006 | < 0.0001 | 0.9108 ± 0.0036 | < 0.0001 |
| 10 | 1.3288 ± 0.0228 | < 0.0001 | 0.8698 ± 0.0224 | < 0.0001 | 0.0462 ± 0.0005 | < 0.0001 | 0.9672 ± 0.0023 | < 0.0001 |
| 30 | 1.1756 ± 0.1838 | 0.34 | 0.7106 ± 0.1276 | 0.02 | 0.0632 ± 0.0009 | < 0.0001 | 1.0472 ± 0.0026 | < 0.0001 |

We determined the fit parameters for simulated catch-slip bond dissociation data (Figure 4 and Tables 3 and S3). The values represent the coefficients ± the standard error of those fits to $(t) = \mathrm{N}e^{-k_{0,c}t}$ for $f = 0$ and Eq. (18a) in the main text for non-zero loading rate with $N = 1000$ and the loading rate entered as fixed parameters because they are known control parameters in an experiment. In the example cases for loading rates 1, 3, and 10 pN/s (Table S3), the fit parameters were significantly different from the value used to stochastically generate the simulated data, *p* values < 0.0001 in all cases (smaller than the double precision data type in MATLAB). However, in the example cases for loading rates 0.3 and 30 pN/s (Table S3), not all the fit parameters were significantly different from the value used to stochastically generate the simulated data, *p* values > 0.05. For low loading rate (0.3 pN/s) the fits to the parameters associated with the slip bond were not significantly different from the simulated values (*p* values = 0.46 and 0.34), and for high loading rate (30 pN/s) the fit to $k_{0,\mathrm{c}}$ was not significantly different from the simulated value (*p* value = 0.34). However, this does not suggest that the fits were better. They were not significantly different from the simulation parameters because the s.e. of the fits were at least two orders of magnitude higher than for intermediate loading cases.

Despite the high standard errors of the fit in the lowest and fastest loading rates, all the example data sets shown in Figures 4b and 4c and Table S3 were from simulations that were well fit. For some of the simulated data sets, we found that the nonlinear least squares function in MATLAB was unable to return a unique, bounded set of fit parameters. To force MATLAB to return values, we set upper limits on the fitting parameters of 10. In cases where the fit values were at or near (> 3) to these upper limits, we discarded the fits from further analysis.

We also determined the fit parameters for simulated catch-slip bond dissociation data with short events, corresponding to those that are indistinguishable from the experimental noise, removed (Table 4) using the fit (nonlinear least squares method) function in MATLAB to $n(t) = n_0 e^{-k_{0,\mathrm{s}}(t-t_0)}$ and Eq. (18b) in the main text. Again, we used the number of "detected" unbinding events in each simulation, $n_0$, $f$, and $t_0 = 50$ ms as fixed parameters in the fitting because the total number of unbinding events detected that were distinguishable from the noise, $n_0$, and the loading rate, $f$, are known with reasonable confidence in a force spectroscopy experiment, i.e., optical tweezers.

We found that the nonlinear least squares function in MATLAB was unable to return a unique, bounded set of fit parameters for more of the simulated data sets when the short events ($t < t_0 = 50$ ms) were discarded, than when those events were not discarded. In addition to setting upper limits on the fitting parameters to 10, we also set lower limits on the fitting parameters to 0. In cases where the fit values were at or near (> 3) to these upper limits or lower (< 0.1 for parameters equal to 1), we discarded the fits from further analysis.
